# Effects of microgravity on the three-dimensional morphology of rhizoids in *Physcomitrium patens*

**DOI:** 10.64898/2026.01.30.702759

**Authors:** Naoki Yagihara, Takahisa Wakabayashi, Ryohei Yamaura, Daisuke Tamaoki, Hiroyuki Kamachi, Daisuke Yamauchi, Yoshinobu Mineyuki, Makoto Hoshino, Kentaro Uesugi, Toru Shimazu, Haruo Kasahara, Motoshi Kamada, Tomomi Suzuki, Yuji Hiwatashi, Yuko Hanba, Atsushi Kume, Tomomichi Fujita, Ichirou Karahara

## Abstract

Rooting systems of plants perceive environmental stimuli and flexibly regulate their growth. Therefore, understanding stimulus perception and response mechanisms is essential for optimizing cultivation. During the transition from aquatic to terrestrial environments, land plants have acquired mechanisms to adapt to gravitational force on land. Thus, elucidating gravity responses of rhizoids in bryophytes, early diverging land plants, provides important insights into how gravity-response mechanisms were established during land plant evolution. Analyzing rhizoid morphology under microgravity, where gravitational effects are largely eliminated, provides an effective approach to examine the gravity-response mechanisms that evolved after terrestrialization. In this study, to elucidate microgravity effects on rhizoid growth of *Physcomitrium patens*, we analyzed 3D datasets obtained by refraction-contrast micro-CT using synchrotron radiation after fixation and embedding of samples from the Space Moss experiment conducted on the International Space Station. Because each CT volume contains numerous rhizoids, we optimized a WEKA-based machine-learning segmentation approach by improving preprocessing, training, and postprocessing steps, resulting in a significantly improved segmentation accuracy. Comparison of 3D morphological indices between manually segmented rhizoids and predicted results supported the validity of the proposed method for morphological analysis. Morphological analyses revealed that, compared with both ground and artificial 1 × g conditions, rhizoid elongation and gravitropic responses were suppressed under microgravity, leading to reduced vertical growth. These findings indicate that gravity plays a fundamental role in rhizoid morphogenesis, and their absence affects growth orientation and elongation. This study provides foundational data for research on the rooting systems of bryophytes in space.

## 1. Introduction

Rooting systems of plants play essential roles in uptake of water and nutrients from the soil, mechanically supporting the plant body, and sensing environmental stimuli to regulate growth in response to external conditions. Understanding how plants perceive and respond to the environmental stimuli in their growing conditions, and how they regulate the growth of underground organs and the aforementioned functions, is a critical issue for optimizing cultivation conditions in a given environment.

Bryophytes are among extant groups of land plants that were the first to colonize terrestrial environments. The rhizoids of bryophytes, which consist of unicellular or multicellular filaments, serve to anchor the aerial parts and to absorb water and, potentially, nutrients, analogously to roots (Jones and Dolan, 2012). As pioneer plants, they colonized the surfaces of rocks and other substrates, promoting the accumulation of organic matter and contributing to soil formation (Jackson, 2015). Rhizoids, which respond to gravity, likely played a key role in this process. Among various environmental cues, gravity plays one of the pivotal roles in determining root orientation and growth regulation. When plants colonized land, having lost buoyancy, plants became directly exposed to a gravitational acceleration of 1 × g on land and therefore evolved mechanisms to cope with gravity. These responses are known as gravity resistance responses (Hoson and Soga, 2003; Soga, 2013). Elucidating the gravity-resistance response in bryophytes offers valuable insight into how land plants first developed gravity-resistance mechanisms and how these systems later evolved into those found in vascular plants. To compare these mechanisms with those of vascular plants, moss serves as a suitable model because it forms a gametophore, consisting of phyllids analogous to leaves and caulids analogous to stems, as well as a rhizoid system analogous to roots, giving it an external morphology resembling a miniature version of vascular plants. Among moss species, *Physcomitrium patens* is a model bryophyte species whose genome has been completely sequenced (Rensing et al., 2008), enabling extensive genetic and cellular studies.

Regarding responses of mosses to the altered gravitational acceleration, Takemura et al. (2017) cultivated *P. patens* under 10 × g for 8 weeks and found that the caulid became shorter and thicker (Takemura et al., 2017), which is similar to the responses of the stems of vascular plants (Nakabayashi et al., 2006; Soga et al., 2006). Takemura et al. (2017) also demonstrated that rhizoid length increased significantly under 10 × g for 8 weeks compared to the 1 × g condition, and that higher gravitational acceleration promoted rhizoid elongation. Similarly, *Arabidopsis thaliana* (L.) Heynh. roots also exhibit enhanced elongation under hypergravity conditions, as observed for *P. patens* rhizoids (Ohara et al., 2025). Regarding the anchorage function of roots, rooting depth and embedded length in soil as major determinants of pull-out resistance (Ennos, 1990; Stokes et al., 1996; Freschet and Roumet, 2017). In contrast, the anchorage function of rhizoids has not been investigated in detail. Moreover, a comprehensive assessment of how altered gravity affects functional aspects of the plants’ rooting structure, such as mechanical support and absorption of nutrient and water, requires observation of its 3D morphology.

Regarding responses of plant root to the altered gravitational vector, roots are known to exhibit positive gravitropism. This response is mediated by amyloplasts functioning as statoliths—dense, starch-filled organelles that sediment within the columella cells of the root cap and trigger gravity perception (Sack et al., 1986; Kiss et al., 1989). Research on the subsequent mechanisms is progressing (Morita, 2010). Generally, rhizoids in bryophytes acquire polarity and grow downward in response to gravitational stimuli (Lobachevska et al., 2022). While amyloplasts are involved in gravity sensing in the protonemata of *Ceratodon purpureus* (Hedw.) Brid. and in the thalli of *Marchantia polymorpha* L. (Kuznetsov et al., 1999; Hashimoto-Sugimoto et al., 2025), in other bryophytes and in rhizoids, the presence of amyloplasts has not been confirmed. This suggests that starch-based sedimentation-type statolith structures may be absent, and the mechanism by which rhizoids perceive and respond to gravity remains unknown.

Recently, we have performed “Space Moss” experiment on the International Space Station to understand effects of microgravity on the growth of *P. patens* (Kume et al., 2021). Although the effects of microgravity should be interpreted in the context of continuous light conditions, under which the plants were grown in this experiment, the influence of microgravity on rhizoid gravitropism can still be observed. The entire set of rhizoids belonging to a single gametophore is called “rhizoid system” (Jang et al., 2011), and the rhizoid systems of *P. patens* obtained from the “Space Moss” experiment were embedded in paraffin and imaged using refraction contrast X-ray micro-computed tomography (µCT) using a synchrotron radiation facility SPring-8 (Yamaura et al., 2022). However, each µCT dataset consisted of approximately 3,000 slices containing numerous rhizoids, making manual segmentation labor-intensive and time-consuming. Automatic segmentation using a machine learning approach was attempted and visualization of the moss rhizoid system in 3D was successful for the first time. Nevertheless, the segmentation accuracy achieved was insufficient for reliable morphological analysis.

In this study, we utilized µCT data of the rhizoid system of *P. patens* obtained from the Space Moss experiment to improve rhizoid segmentation performance. We examined machine learning parameters as well as preprocessing and postprocessing strategies, and also evaluated segmentation performance in 3D. We aimed to quantitatively analyze the morphological responses of *P. patens* rhizoids under microgravity, space 1 × g, and ground 1 × g control conditions. Through this analysis, we aimed to elucidate the effects of microgravity on rhizoid growth and 3D-morphology.

## 2. Materials and methods

### 2.1 Plant materials and growth conditions

Plants of a moss species *Physcomitrium patens* (Hedw.) Mitt. were used in the present study. The specimens were obtained from the Space Moss experiments (Run 1, 2) conducted between 2019 and 2020. The cultivation conditions for the Space Moss experiments were as follows. Forty-eight gametophores were planted on agar slabs (W×D×H= 54 mm × 44 mm × 25 mm) supplemented with BCD medium (Nishiyama et al., 2000). These slabs were placed inside polycarbonate growth chambers with external dimensions of 60 mm × 50 mm × 60 mm. The chambers were housed in Plant Experimental Units (PEUs), which were specifically designed for plant research aboard the International Space Station (ISS) (Karahara et al., 2020).

Prior to the initiation of the experiments, the plants were kept refrigerated. The PEUs were installed either in the microgravity compartment or on the centrifuge of the Cell Biology Experimental Facility (CBEF) within the ISS (Ishioka et al., 2004; Yano et al., 2013), where artificial gravity equivalent to 1 × g was applied (Yashiro et al., 2020). This condition is referred to as "Space 1 × g." For ground-based control experiments, equivalent PEUs were installed in the CBEF located at the Tsukuba Space Center.

The plants received illumination from above using a matrix of light-emitting diodes (LEDs) (Yano et al., 2013), providing a light intensity of 30 µmol m⁻² s⁻¹ at the center of the bottom surface of the growth chamber. The cultivation period lasted for 25 to 26 days at a constant temperature of 25 °C. The Run 1 experiment was conducted between July and August 2019, and Run 2 was carried out from December 2019 to January 2020. In this study, a rhizoid system from one individual gametophore was randomly selected from each of the following growth chambers used in the Space Moss experiments: Ground 1 × g conditions (#27, #82, #84), Space 1 × g conditions (#003, #004, #103), and Space µ×g conditions (#002, #101, #B101). These samples were designated as Ground 1 × g_27, 82, 84; Space 1 × g_003, 004, 103; and Space µ × g_002, 101, B101, respectively.

### 2.2 Chemical fixation and embedding of rhizoid system specimens

*P. patens* cultivated in orbit during the Run 1 and Run 2 experiments were chemically fixed aboard the ISS together with the agar medium. The fixation solution contained 2.5 % (v/v) glutaraldehyde (Polysciences, Inc., Warrington, PA, USA) in 100 mM sodium phosphate buffer and adjusted at pH 7.2. Following fixation, the samples were stored under refrigerated conditions at 2 to 4 °C until the embedding process was initiated on the ground (Fig. 1a).

**Fig 1.**
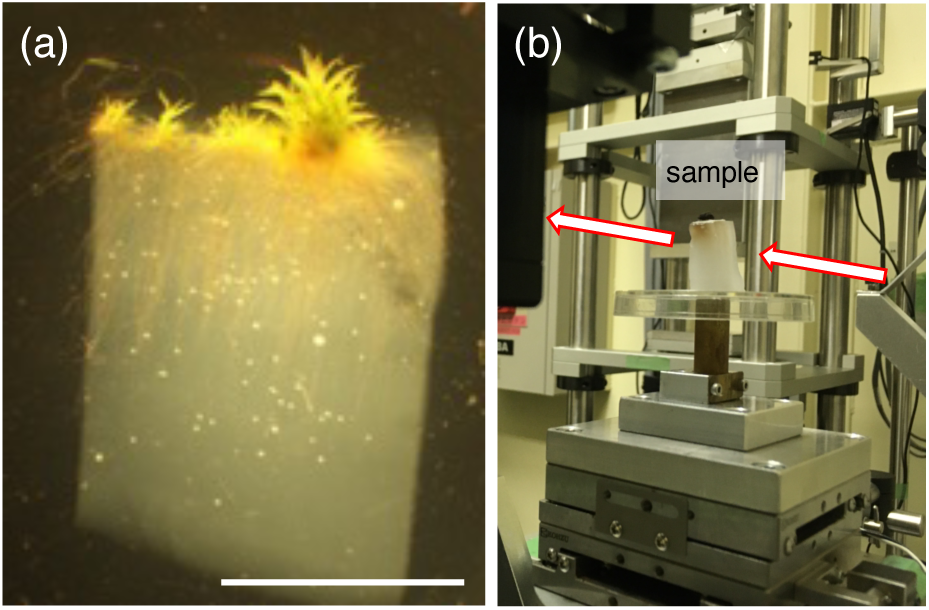
Specimen preparation and X-ray micro-CT scanning. (a) Lateral views of gametophores and rhizoids of P. patens grown on/in agar media observed under a dissecting microscope. Specimen example: Ground 1 × g_27. Scale bars = 10 mm. (b) Experimental setup at the beamline BL20B2. Arrows indicate the X-ray path. Rhizoid system embedded in paraffin was placed on a rotation stage.

The fixed samples were primarily embedded in paraffin for subsequent µCT analysis. Specimens obtained from Run 1 were dehydrated through a graded ethanol series (70 %, 90 %, 99.9 %, and 100 % [v/v]), followed by xylene treatment, all performed in glass vials placed on a rotary shaker set at 80 rpm. After dehydration, the samples were infiltrated with paraffin (Paraplast Plus, Sigma-Aldrich Japan, Tokyo) at 60 °C for a period of three days.

### 2.3 Refraction-contrast X-ray micro-CT and tomographic reconstruction

Refraction contrast micro-CT was performed at Experimental Hutch 1 of beamline BL20B2 at the SPring-8 synchrotron radiation facility operated by the Japan Synchrotron Radiation Research Institute (JASRI), following the procedures described previously (Kurogane et al., 2021; Yamaura et al., 2022). The experimental setup is illustrated in Fig. 1b, and a summary of the imaging protocol is provided below. X-rays were set to an energy level of 15 keV. The projected images on the scintillation screen were continuously captured using a CMOS detector (ORCA-Flash 4.0, Hamamatsu Photonics, Hamamatsu, Japan). Each image measured 2048 × 2048 pixels in resolution, corresponding to a physical field of view of approximately 5 × 5 mm. A total of 900 projection images were recorded over a 180° rotational range.

Due to the spatial distribution of rhizoids extending more than 5 mm in depth from the base of the gametophore, and because most rhizoids in the specimens used in this study were contained within 10 mm, two separate scans were performed: one scan covering the area from the base to a depth of 5 mm, and the other from 5 mm to 10 mm. For the specimens used in this study, the effective voxel size of the reconstructed 3D volumes was estimated to be 2.75 µm. Tomographic reconstruction was carried out using a convolution back-projection method, and image processing was performed with the Chesler filter available in the SP-µCT software suite (http://www-bl20.spring8.or.jp/xct/) (Kurogane et al., 2021).

### 2.4 Image preprocessing

Because volume data of the rhizoid system were acquired from both proximal and distal regions, corresponding to the upper and lower on the ground regions, background pixel values were normalized. After converting 16-bit grayscale images into 8-bit, the background pixel values outside the field of view were set to 0, and the mode pixel value within the circular field of view was adjusted to 15. Maximum pixel values were set to 80-100. After that, to enable machine learning to work effectively on both volumes, a "3D Median" filter (X-Radius: 2, Y-Radius: 2, Z-Radius: 2) and an "Unsharp Mask" filter (Radius: 27.0, Ratio: 0.3) were applied using Fiji software package (https://imagej.net/software/fiji/) (Schindelin et al., 2012), in order to reduce background noise and enhance the contours of rhizoids. After this operation, the region containing only paraffin without agar located in the most proximal region of the proximal volume as well as the region containing gametophore were excluded. A duplicated region was present at the most distal region of the proximal volume and the most proximal region of the distal volume. To eliminate the duplicated (overlapping) region between the proximal and distal volumes, we removed the most proximal approximately 200 slices of the distal volume, where noise was abundant. Approximately 900 slices in the proximal volume and 1800 slices in the distal volume were combined, and the combined volume of about 2,700 slices in Z-axis was obtained.

### 2.5 Automatic segmentation

For the automatic segmentation of rhizoids in the tomogram, the plugin Trainable Weka Segmentation 3D (TWS 3D) (Arganda-Carreras et al., 2017) of Fiji software package was used. Because the number of slices that could be labeled at one time was limited to about 200 due to the specifications of the desktop computer (Intel Core i9 CPU, 256 GB RAM) used in the present study, the combined volume was divided every 200 slices, i.e., 200 pixels in thickness, and machine learning models were created. In the cross-sectional tomographic slices, rhizoids were observed as three types (Fig. 2), i.e., Type A, in which a rhizoid was properly embedded in paraffin and delineated by a cell wall (Fig. 2a), Type B, in which a rhizoid was flattened during embedding (Fig. 2b), and Type C, in which a void space is formed after a rhizoid is collapsed due to poor infiltration of paraffin (Fig. 2c). These are the same types as those categorized by Yamaura et al. (2022). Therefore, three types of annotations were made: the cell walls of rhizoids represented by Type A and B objects as "ab", void space of Type C (black area surrounded by a bright halo) objects as "c", and the background area where rhizoids do not exist or outside the field of view as "bg". For the filters used for feature extraction during training, Mean, which captures the overall brightness and darkness in the image, Variance, which captures the texture and boundary complexity of the image, and Hessian, which detects curvature, were adopted. The parameter Max Sigma, which is the upper limit of the filter scale, was set to 4.0, and Fast Random Forest classifier was used. For the development of the classifier, a substack comprising 200 slices was extracted from the position located at 25 % from the top in depth (z-axis) direction, and 5 to 7 annotations for each of the three classes were made. Subsequently, the initial segmentation results generated by the classifier were manually refined by correcting approximately five regions per class. The classifier was then retrained using the modified data. Following this refinement, the updated classifier was applied to a segmented substack comprising 200 slices extracted from the position located at 50 % from the top of the volume in z-axis. Within this segmented substack, approximately five regions per class were manually corrected again. An identical refinement process was conducted for a substack extracted from the position located at the 75 % from the top of the volume in z-axis. These three refinement steps performed at 25 %, 50 %, and 75 % in z-axis were defined as a single correction cycle. In the present study, a total of one to five such refinement cycles were conducted. Finally, all substacks were labeled, and by reintegrating the binarized 200-slice substacks, a binarized volume in which rhizoids were labeled was obtained.

**Fig. 2.**
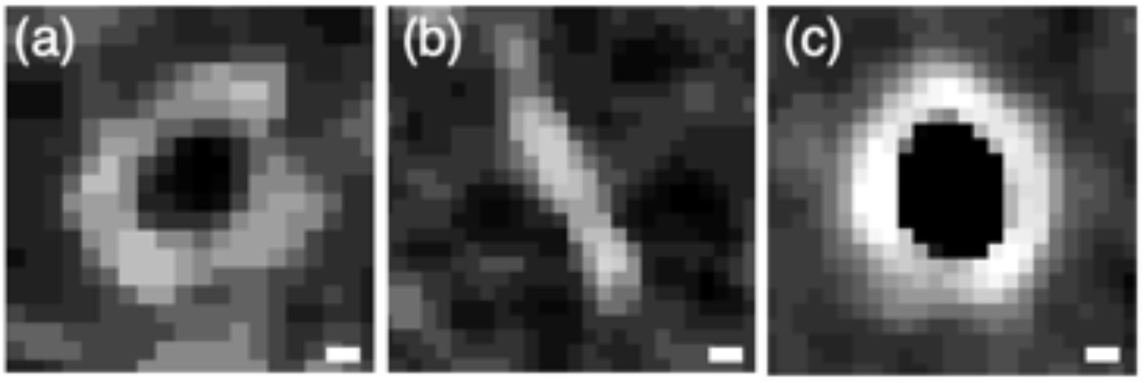
Rhizoid structures observed in CT images after image preprocessing. (a) Type A, rhizoid properly embedded in paraffin and delineated by a cell wall. (b) Type B, rhizoid collapsed and flattened during embedding, resulting in a compressed, linear appearance. (c) Type C, void space formed at a rhizoid location due to poor infiltration of paraffin to the rhizoid, showing the void space as black area surrounded by a bright halo artefactually generated during reconstruction. Specimen example: Ground 1 × g_27. Scale bars = 10 µm.

### 2.6 Image postprocessing

On the binarized image, for the labeling of Type A, a cell wall area was labelled as an incomplete or a complete ring-shaped area without being labeled inside (Fig. 3). Therefore, because of incompleteness of ring-shaped area, a closing process was applied to join adjacent objects into a complete ring shape, and then labeling of the inside was performed using Fill Holes, which labels the inside of closed structures. After that, to reduce noise, a 3D Mean filter was applied with parameters X-Radius: 2, Y-Radius: 2, Z-Radius: 2. After processing with the 3D Mean filter, the image becomes grayscale and is no longer a binary image. Therefore, binarization was performed again by applying a threshold at brightness value of 127. Finally, prominent artifactual objects that should be classified as background, such as void space caused by air bubbles, were manually removed.

**Fig. 3.**
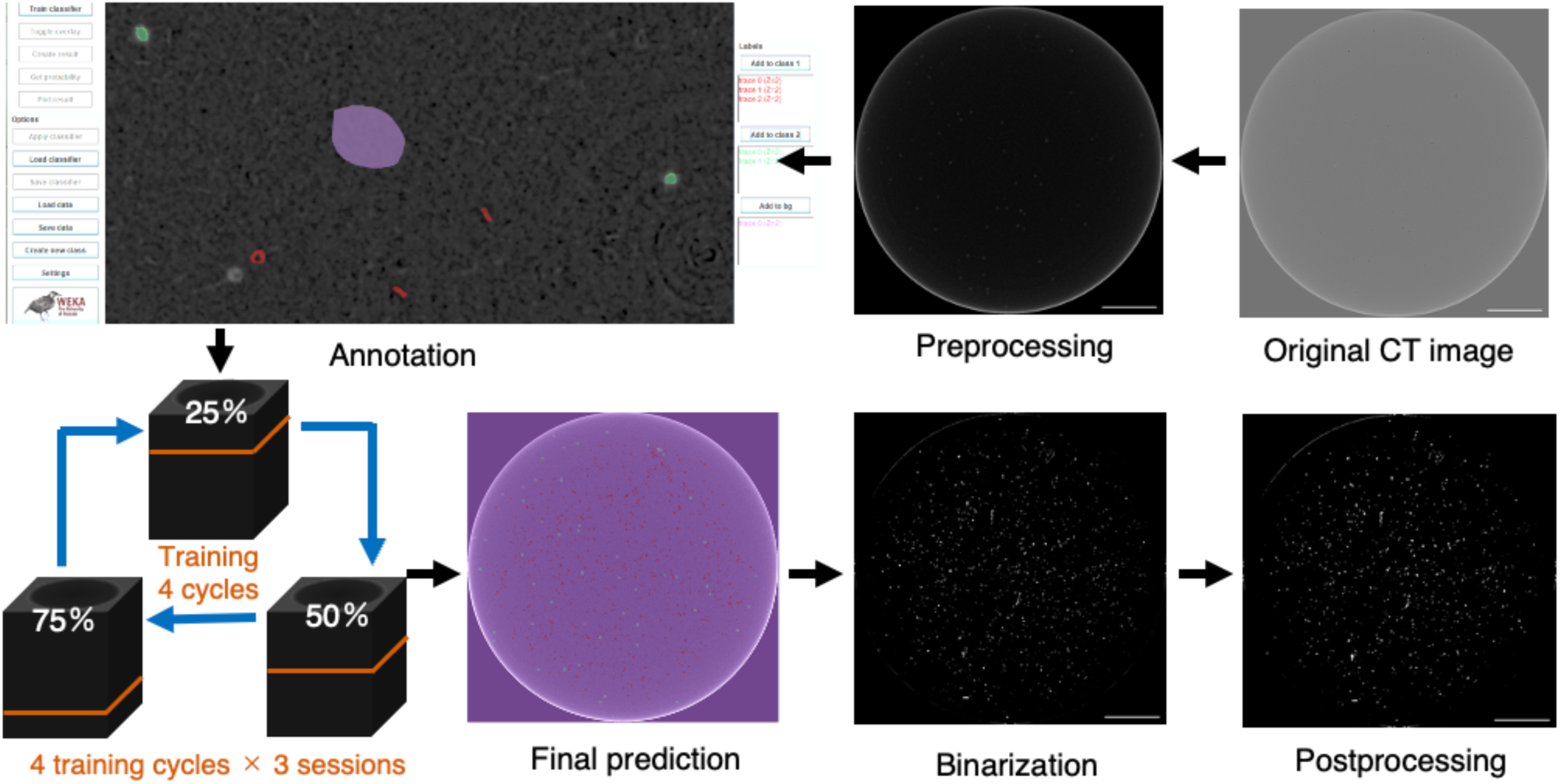
Workflow of automatic segmentation in the present method. The present method is an improved version of the previous approach based on TWS 3D. During both annotation and segmentation, Types A and B were classified into the red-labeled region, Type C into the green-labeled region, and the background into the purple-labeled region. Specimen example: Ground 1 × g_27. Scale bars = 1 mm.

### 2.7 Comparison with our previous method

To compare the results obtained using the present method with those from our previous method (Yamaura et al., 2022), we also performed automatic segmentation using the previous method, which is briefly described as follows.

In the previous method, no image preprocessing was performed. Classifiers were trained separately for the proximal and distal volumes without merging them, and segmentation of rhizoids was carried out independently for each. The proximal volume was divided into substacks, each consisting of 100 slices. A representative substack exhibiting typical rhizoid distribution was selected for classifier training. In this method, Types A, B, and C were treated as a single class, with the background defined as the second class, resulting in a two-class classification scheme. During training, the default settings for filters were used without modification, specifically employing the Mean and Variance filters. The Max Sigma parameter was kept at the default value of 8.0. As in the present method, the classifier used was Fast Random Forest. From a substack consisting of 100 slices, approximately five instances of each of the three Types were annotated every 20 slices, and several background regions were selected and annotated across the entire volume to construct the classifier. For classifier refinement, the same substack used in the initial training was employed. In each refinement cycle, approximately five corrections were made for each of the three Types and for the background. This refinement process was repeated five times. The final corrected classifier was then used to perform segmentation across all substacks. In post-processing, Closing was applied with parameters X-Radius: 1, Y-Radius: 1, and Z-Radius: 1, followed by the Fill Holes operation. However, no additional filtering was performed. The same procedures were conducted for the distal volume, and the segmentation images from the proximal and distal volumes were subsequently merged to obtain the complete rhizoid segmentation.

### 2.8 Assessment of automatic segmentation performance

To assess the performance of automatic segmentation, a comparison was made between images derived from the volume after post-processing, i.e., predicted images, and those in which rhizoids were manually labeled from CT images, i.e., ground truth images. At six slice positions corresponding to 14.2, 28.5, 42.8, 57.1, 71.4, and 85.7 % of the depth from the distal end of the region derived from the distal volume (excluding the proximal end corresponding to 100 %), three non-overlapping square regions of 256 pixels per side, with different planar coordinates and excluding the shoot (gametophore) if present, were extracted for each position and annotated, yielding 18 ground truth images in total. Also, since in some samples the number of slices in the proximal volume was approximately half that of the distal volume, three regions were similarly extracted at 25, 50, and 75 % depth from the region derived from the proximal volume, yielding 9 ground truth images in total.

For each assessment square region of 256 pixels per side, a confusion matrix was created, and from the values of the confusion matrix, the pixel accuracy (PA), F1-score (F1) and Intersection Over Union (IoU) between the ground truth and predicted images were calculated (Supplementary Fig. 1). The values of the confusion matrix and the calculation of each index were performed using a program written in Python. In addition to these metrics, precision and recall were used as evaluation indices to compare the false positive and false negative ratios of the extracted regions. Precision decreases with an increase in over detections, and recall decreases with an increase in missed detections. The F1 score is high when precision and recall are simultaneously high and balanced.

### 2.9 Morphological Analysis

For the post-processed volume, objects with 10 voxels or less were removed for the removal of noises. Next, the areas of the object were calculated for each slice, and objects corresponding to parts of the shoot were removed. The shoot object was removed based on 2D criteria to minimize the removal of rhizoid objects that emerged directly from the shoot. Then the morphological analysis was performed. In the depth direction of the volume, the range from 1–33.3 % from the top was defined as the proximal part, 33.4–66.6% as the middle part, and 66.7– 100 % as the distal part, and for each as well as for the entire, the following parameters were analyzed: total rhizoid volume, total rhizoid surface area, total rhizoid length, average rhizoid diameter, and rhizoid growth angle.

#### 2.9.1 Total volume and total surface area of rhizoids

Using “AnalyzeRegion3D,” a plugin of Fiji, total volume and total surface area per 1 mm³ were calculated.

#### 2.9.2 Total length of rhizoids

In the measurement of rhizoid length, objects were skeletonized using Skeletonize (2D/3D) plugins of Fiji software and represented as medial axis of 1-pixel width. Using “AnalyzeSkeleton,” length of continuous line called ‘branch’ was measured as rhizoid length, and the sum of the lengths of all rhizoids was taken as the total rhizoid length. Then, the total rhizoid length per 1 mm³ was calculated.

#### 2.9.3 Average rhizoid diameter

Using the “Analyze Particles” function in Fiji, the minimum diameter of each object in a single slice was calculated across all slices in the volume. In this study, since rhizoids grew in various directions, the minimum diameter was defined as the rhizoid diameter because the maximum diameter is influenced by the growth direction.

#### 2.9.4 Rhizoid growth angle

For growth angle analysis, rhizoids with a tubular shape were selected from the volume. For shape discrimination of predicted objects, “vesselness score” (Phalempin et al., 2021) was used. The shape of the object was approximated as an ellipsoid, and vesselness score (v) was calculated. The more the approximated ellipsoid of the object is elongated and tubular, the closer v becomes to 1. In this study, to use only objects with v larger than 0.8 for angle quantification, other objects (referred to as non-tubular objects) were removed. However, many of the rhizoids having branching and/or contacting parts were removed by doing this object discrimination because these parts have v scores smaller than 0.8. Therefore, before doing this object discrimination, blank slices were intentionally inserted every 400 slices into the stack to artificially split rhizoids, thereby limiting the size of non-tubular objects to less than 400 slices (Supplementary Fig. 2). Using the fiber analysis software open_iA (Fröhler et al., 2019), the angle θ (0° ≦ θ ≦ 90°) formed with the Z-axis by the line segment connecting the both ends of the object was calculated from the volume filtered by vesselness score. Rhizoid growth angle per object was first measured. However, rhizoids were fragmented in the volume data. In samples where degree of fragmentation is high, angles of short objects are overestimated if usual average, i.e., arithmetic mean, is calculated. Therefore, the sum of angle values weighted by rhizoid length per object, i.e., multiplication of angle by length, was divided by the total rhizoid length, and angle of the entire rhizoid system weighted by rhizoid length was obtained, which is referred to as length-weighted angle value.

## 3. Results

### 3.1 Optimizing conditions of training cycle and Closing

To improve the segmentation conditions, we first investigated the optimal number of training cycles and the appropriate parameters for post-processing using the closing operation. While our previous method performed training within the same substack (Yamaura et al., 2022), the present method aimed to enhance generalizability of the classifier by training sequentially within the substacks located at 25 %, 50 %, and 75 % regions from the top of the volume. One training cycle was composed of the training at three locations (25 %, 50 %, and 75 %), and we examined how many such cycles would be the most effective.

Although the F1 score generally increased with additional training cycles, some samples, such as those from the Space 1 × g group, showed decreased F1 scores when trained for four or five cycles (Supplementary Fig. 3). This is considered to be the result of overfitting. It is likely that the model became overly adapted to the characteristics of the substack used for training, leading to reduced generalization performance and a consequent drop in the F1 score when segmenting the entire volume. The median number of cycles at which the highest F1 score was achieved was 4. Based on these results, we determined that four training cycles were optimal.

Next, we examined the optimal parameters for the closing operation to form complete ring-shaped rhizoid structures of Type A, while minimizing undesired connections between neighboring predicted objects. In the previous method, closing was applied with the following parameters set, X, Y and Z radii are 1, 1 and 1 pixel, respectively (referred to as Closing 1). In the present study, we tested three different closing conditions as follows: all the X, Y and Z radii set to 2 (Closing 2), 3 (Closing 3), in addition to 1 (Closing 1), respectively. For each condition, the binary volume was processed with the corresponding closing operation, followed by application of Fill Holes and a 3D mean filter (X, Y, and Z radii are 2, 2 and 2 pixels, respectively). F1 score and IoU were calculated based on the resulting volumes. To ensure a fair comparison of closing effects, manually corrected volumes were not used in this evaluation.

As a result, among all 9 samples throughout the different gravity conditions, the highest F1 score was achieved in 2 samples with Closing 1, in 6 samples with Closing 2, and in 1 sample with Closing 3 (Supplementary Fig. 4a). A similar trend was observed for IoU values (Supplementary Fig. 4b). The rhizoid objects of Type A sometimes failed to form complete ring-like structures with Closing 1, while excessive merging of neighboring objects was observed with Closing 3. In contrast, Closing 2 resulted in neither incomplete closure of rhizoid structures nor excessive merging of adjacent objects (Supplementary Fig. 5). These results indicate that Closing 2 provides the most appropriate balance between object closure and suppression of over-merging, thereby effectively improving segmentation performance by preserving rhizoid morphology. Therefore, Closing 2 was adopted as the standard parameter for postprocessing in rhizoid object segmentation in this study, based on its superior performance by well-preserving rhizoid morphology.

However, Closing 1 yielded the highest F1 score and IoU in Ground 1 × g_84 and Space 1 × g_103, while Closing 3 produced the best results in Space µ × g_002. The 2 samples, Ground 1 × g_84 and Space 1 × g_103, included regions in which rhizoid objects were densely distributed. In such regions, the short distances between adjacent objects likely led to inappropriate merging under Closing 2 or 3, resulting in lower segmentation performance (Supplementary Fig. 6). Therefore, Closing 1, which did not cause excessive merging of objects, achieved the highest F1 score and IoU. In contrast, Space µ×g_002 exhibited a relatively low density of rhizoid objects, with individual objects being spatially well separated. As a result, even when Closing 3 was applied, round objects maintained their shape, and the structural completion it provided likely contributed to improved segmentation performance (Supplementary Fig. 7).

Distribution of the number of objects along the depth of the volume was analyzed (Supplementary Fig. 8). Artifactual peaks were found either in the region near the junction between the proximal and distal volumes, likely caused by noise. These noises were effectively removed either by the normalization of the pixel values in the preprocessing, optimization of training cycles, or usage of filters in the postprocessing.

### 3.2 Comparison of segmentation performance between the previous method and the present method

We compared the segmentation performance of the previous method, which involved no preprocessing, training a classifier five times on the same set of 100 slices, and applying Closing 1 and Fill Holes as postprocessing, with that of the present method, which included preprocessing such as brightness adjustment and filtering, training a classifier four times each on three evenly spaced regions of 200 slices (i.e., 4 training cycles × 3 sessions), and applying Closing 2, Fill Holes, and additional filtering as postprocessing. Among the nine samples, the F1 score significantly increased in eight samples with the present method compared to the previous method. Even in Ground 1×g_82, where no statistically significant difference was observed, both the mean and median F1 scores improved (Supplementary Fig. 9). Furthermore, in Space µ × g_002, which initially had the lowest F1 score, the present method raised the median score to 0.637 (Supplementary Table 1). In addition, the interquartile range of F1 scores, represented by the box length in box plots, was narrower with the present method than with the previous method in most samples, indicating reduced variability in performance. This suggests that the prediction performance became more stable across test images, and the generalization ability of the classifier was improved. These results demonstrate that the present method not only improved overall F1 scores but also enhanced stability compared to the previous method, making it an effective approach for rhizoid segmentation.

However, as observed in the training cycle optimization, the Space µ × g samples exhibited large variation in F1 scores, indicating that classifier performance was inconsistent across images. This can be attributed to substantial variation in rhizoid (foreground) density depending on the depth at which the ground truth images were acquired, which was more pronounced in Space µ×g than in other gravity conditions. As a result, test images showed uneven rhizoid densities—high-density images yielded better predictions, whereas low-density images led to poorer detection performance, causing greater variability in F1 scores. Because the classifier had not sufficiently learned to generalize across such differences in density during training, it struggled to maintain stable segmentation performance when applied to test images with varying densities.

Finally, regarding the median values of F1 score, IoU, precision, and recall across samples, the present method outperformed the previous method in all metrics except for recall in Space 1 × g_003 and 004 (Supplementary Table 1). In these two samples, precision was notably higher, while recall declined. This is likely due to enhanced suppression of false positives during the training phase of the present method, which led to fewer over-predictions of rhizoid objects. In other words, although prediction reliability (precision) improved, recall decreased due to some rhizoid objects not being detected—a typical trade-off between the two metrics. Nevertheless, in terms of the F1 score, which integrates both metrics, the gain in precision outweighed the drop in recall, resulting in an overall improvement in segmentation performance.

### 3.3 Three-dimensional assessment of segmentation performance

Two-dimensional assessment of segmentation performance using ground truth images demonstrated that the segmentation performance of the present method was superior to that of the previous method. To further assess the validity of the present method in the morphological analysis of rhizoid structures, a three-dimensional evaluation was conducted. Specifically, morphological parameters were extracted from annotated volumes and their corresponding predicted volumes, and the results were statistically compared to evaluate the validity of morphological analysis based on the predicted volumes. As a representative sample, we selected Ground 1 × g_82, which exhibited an F1 score equal to the median among the Ground 1 × g samples processed using the present method. Within the 3D volume of this sample, three distinct 256 × 256 pixels square regions were selected from different lateral positions between the 201st and 400th slices from the proximal end. Each region was manually annotated, resulting in three annotated volumes comprising 200 slices each (256 × 256 × 200 pixels) (Fig. 4a).

**Fig. 4.**
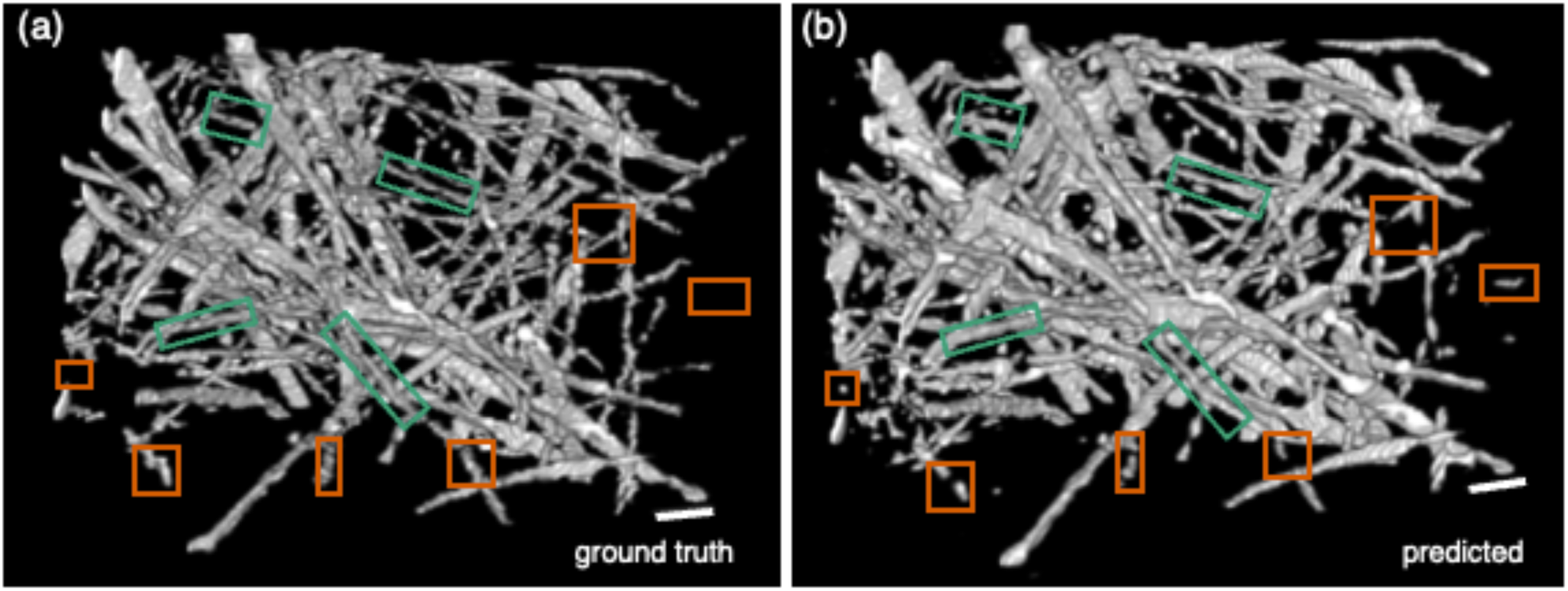
3D models of the annotated ground truth volume (a) and the predicted volume (b). The Orange boxes indicate regions in the predicted volume where rhizoid fragmentation and noise labeling were observed, while the green boxes highlight areas where rhizoids were noticeably over-labeled with increased thickness. Ground 1 × g_82. Scale bars = 0.1 mm.

A comparison between the annotated volumes and the predicted volumes revealed the following two notable issues (Fig. 4). First, the predicted volumes exhibited a high degree of rhizoid fragmentation and excessive labeling of background noise (Fig. 4b). In particular, numerous thin and short fragmented labels were distributed throughout the volume, which likely resulted from the classifier’s failure to correctly distinguish rhizoids with low contrast from the background, or to exclude noises with intensity values similar to the rhizoids. As a consequence of this rhizoid disconnection and over-detection of noise, the number of labeled objects was significantly higher in the predicted volumes than in the annotations (Fig. 5f). Second, the predicted rhizoids tended to appear slightly thicker and exhibited expanded contours compared to the ground truth (Fig. 4b). This phenomenon is primarily attributed to the smoothing effect of the mean filter applied during postprocessing, which may have blurred the boundary regions and caused expansion of the labeled structures. On the other hand, the mean diameter was significantly lower in the predicted volume than in the ground truth, giving the impression that the rhizoids were labeled more narrowly (Fig. 5e). This apparent contradiction is likely due to the presence of a large number of fine noise labels in the predicted volume, which reduced the overall average diameter. These observations suggest that the segmentation method could be further improved.

**Fig. 5.**
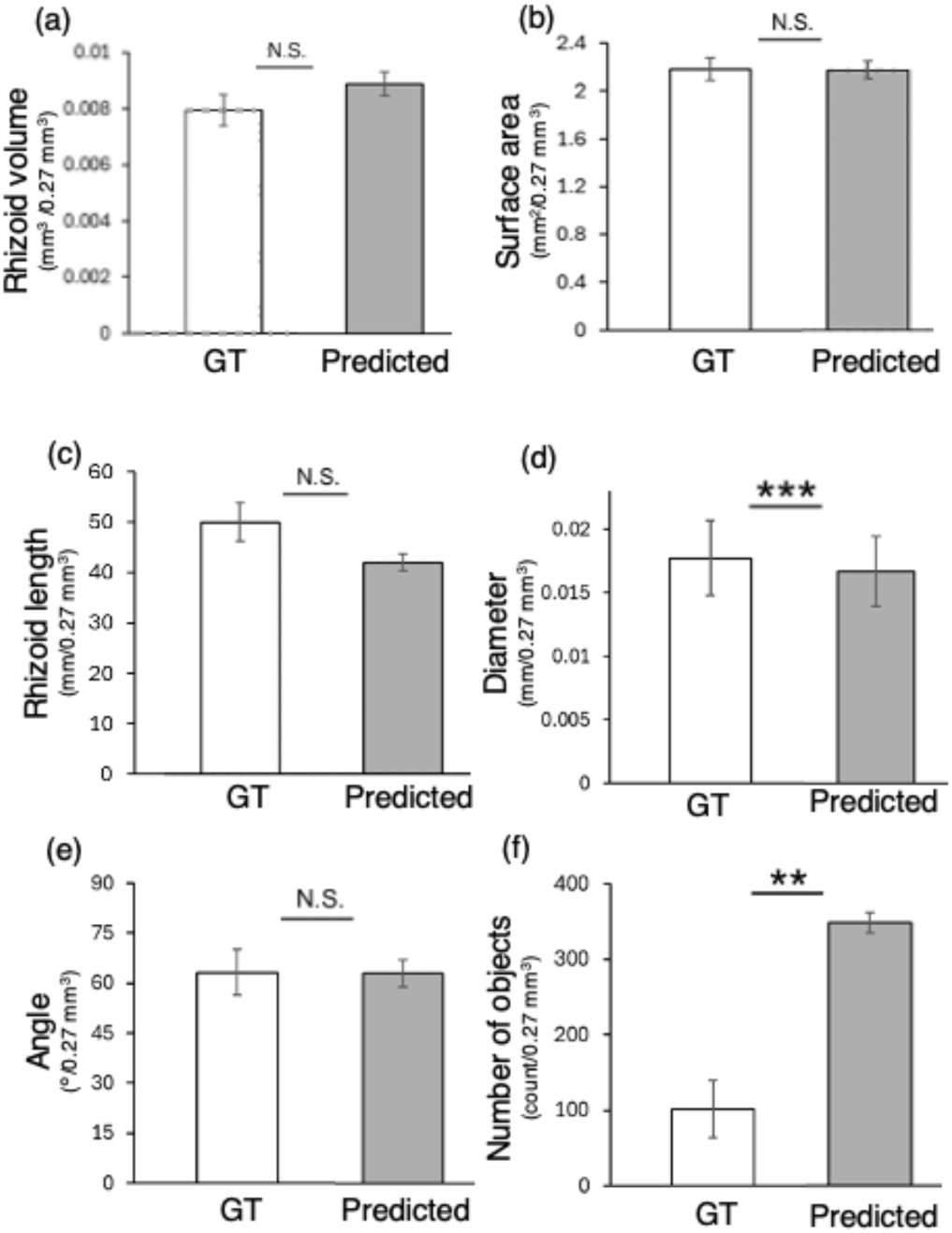
Comparison of morphological parameters of the rhizoid between the ground truth (GT) and predicted volumes. (a) Total rhizoid volume. (b) Total rhizoid surface area. (c) Total rhizoid length. (d) Length-weighted rhizoid growth angle. (e) Rhizoid diameter. (f) Number of objects. Data represent mean ± SE (a, b, c, f) or Least-squares mean (LS mean) ± SE (d, e). n = 3. (a)〜(c), (f) Student’s *t*-test was performed. (d), (e) EMS-base ANOVA. Fixed effect, ground truth vs predicted. Random effects, local extracted volume. (d) *MS* = 559.875, *F* (1, 1502) = 0.1451, *P* = 0.7033. (e) *MS* = 0.00011, *F* (1, 38988) = 77.661, *P* < 0.0001.: N.S., not significant (*P* ≥ 0.05); **, *P* < 0.01; ***, *P* < 0.001. Ground 1 × g_82.

On the other hand, no statistically significant differences were observed for the following morphological metrics: total rhizoid volume, total surface area, total rhizoid length, and raw average rhizoid growth angle between the ground truth and predicted volumes (Fig. 5a-e). Therefore, the present method is demonstrated to be well-suited for three-dimensional quantitative analysis of rhizoid structures, offering both automation and high accuracy compared to our previous method, though caution is required when comparing parameters other than these specific ones.

### 3.4 Effects of different gravity conditions on the morphological parameters of the rhizoid

Finally, morphological analysis was conducted to examine the effects of microgravity on rhizoid development in the three different parts in depth, i.e., proximal, middle, and distal regions from the gametophore position. A 2-D analysis of the distribution of object counts along the depth of the volume revealed that the density of rhizoids decreased from the proximal to the distal region (Supplementary Fig. 8). This trend was also confirmed by the 3-D analysis (Fig. 6). Among the morphological parameters, in particular, total rhizoid volume (Fig. 6a), total surface area (Fig. 6b), and total rhizoid length (Fig. 6c) were reduced in the Space µ × g group, which is obvious in the proximal regions, indicating an inhibition of rhizoid elongation or branching under microgravity conditions. It is assumed that microgravity effects become less evident in the middle and distal regions is likely due to the reduced number of rhizoids toward the distal end, which makes these regions more susceptible to noise.

**Fig. 6.**
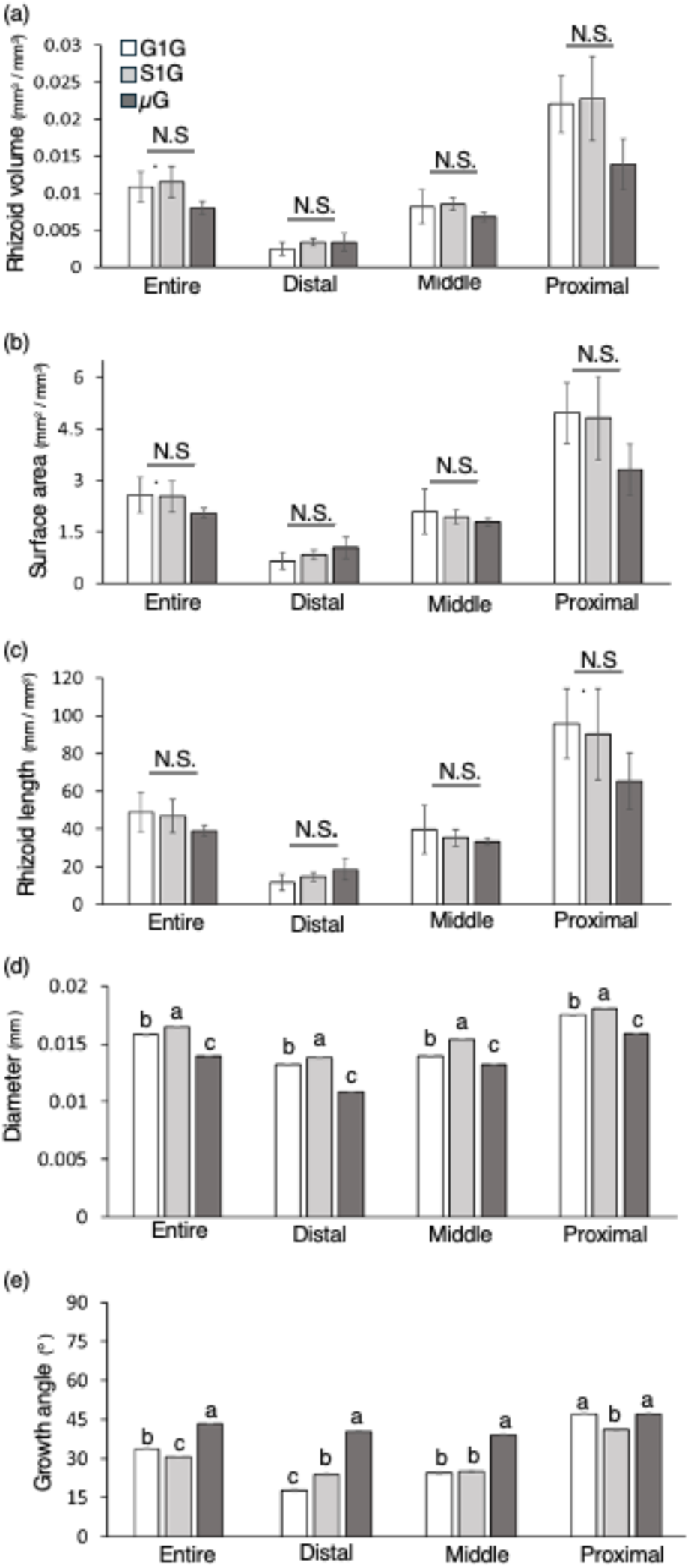
Effects of different gravity conditions on the rhizoid morphological parameters in the entire, proximal, middle, and distal regions. (a) Total rhizoid volume. (b) Total rhizoid surface area. (c) Total rhizoid length. (d) Rhizoid diameter. (e) Raw rhizoid growth angle. Data represent mean ± SE (a-c) or Least-squares mean (LS mean) ± SE (d, e). (a-c) n = 3. (d) n = 6763500 (Ground 1 × g), 6023711 (Space 1 × g), 6049255 (Space µ × g). (e) n = 43905 (Ground 1 × g), 34154 (Space 1 × g), 31291 (Space µ × g). (a-c) One-way ANOVA. N.S., not significant. (d, e) Nested ANOVA (d), Length-weighted nested ANOVA (e). Sample nested within gravity conditions. (d) *F* (2, 2) = 84958.11, *P* < 0.0001. (e) *F* (2, 2) = 2300.937, *P* < 0.0001. (d, e) Different alphabetical letters indicate significant differences in the means assessed by the Tukey–Kramer HSD test (*P* < 0.05) followed by nested ANOVA.

Regarding rhizoid diameter, the dataset comprised measurements from all slices; therefore, these records were not linked to length measurements. Consequently, length-based weighting could not be applied. As a result, the diameters of longer rhizoid may have been overestimated. Under such conditions, however, all regions exhibited statistically significant differences across gravity conditions, with the Space µ × g group showing the smallest mean value and the Space 1 × g group the highest (Fig. 6d).

Regarding the angle values, the columns in Fig. 6e represent the raw rhizoid growth angle, whereas the statistical results correspond to the length-weighted angle analysis. When comparing the space conditions, the mean length-weighted angle was significantly larger in the Space µ × g group than in the Space 1 × g group. In addition, except in the proximal region, the raw rhizoid growth angle showed the highest mean value in the Space µ × g group (Fig. 6e). In both the Ground 1 × g and Space 1 × g conditions, the rhizoids predominantly grew in the distal direction, as shown in the overall 3D model view (Fig. 7) and the polar histogram of raw angle values (Fig. 8).

**Fig. 7.**
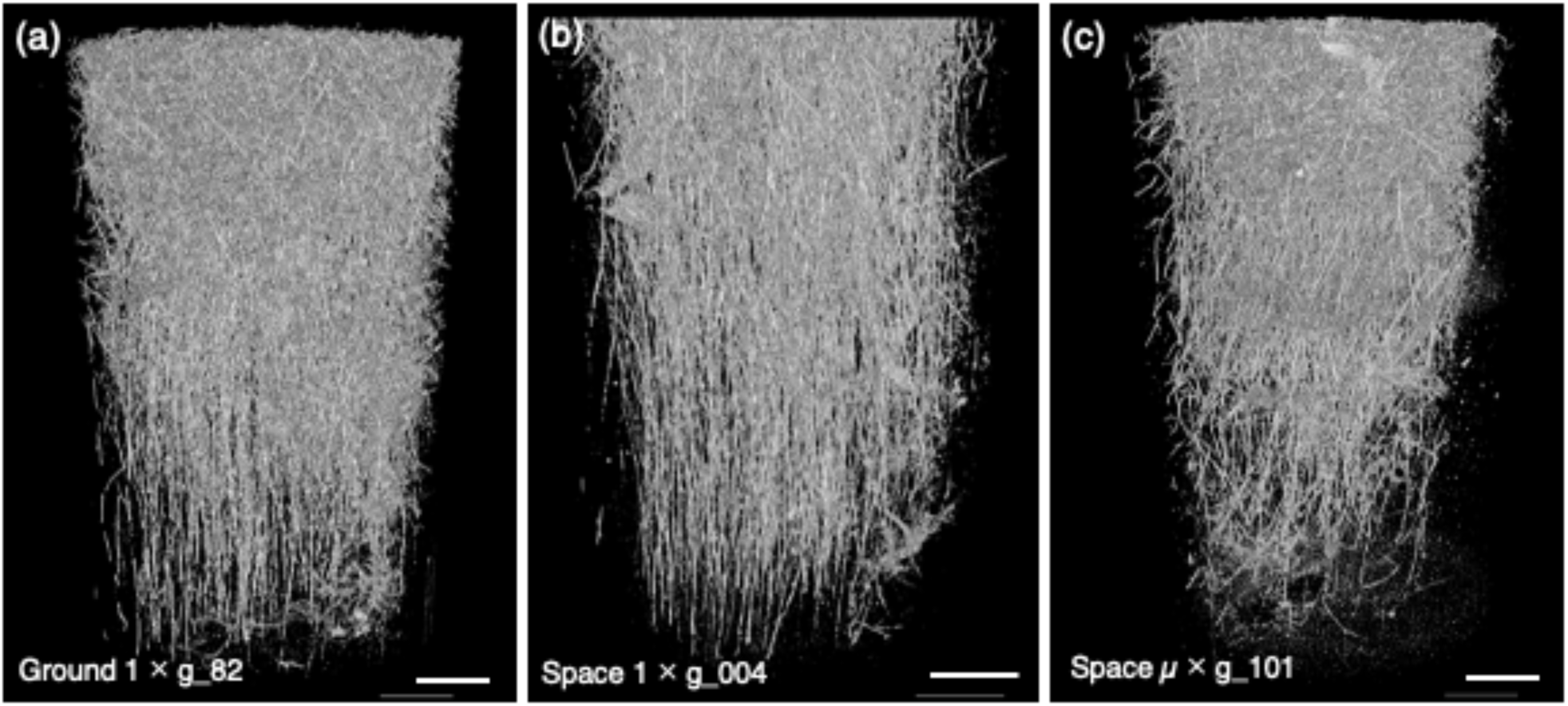
3D models of the rhizoid system architecture grown under Ground 1 × g (a), Space 1 × g (b), or Space µ × g (c).

**Fig. 8.**
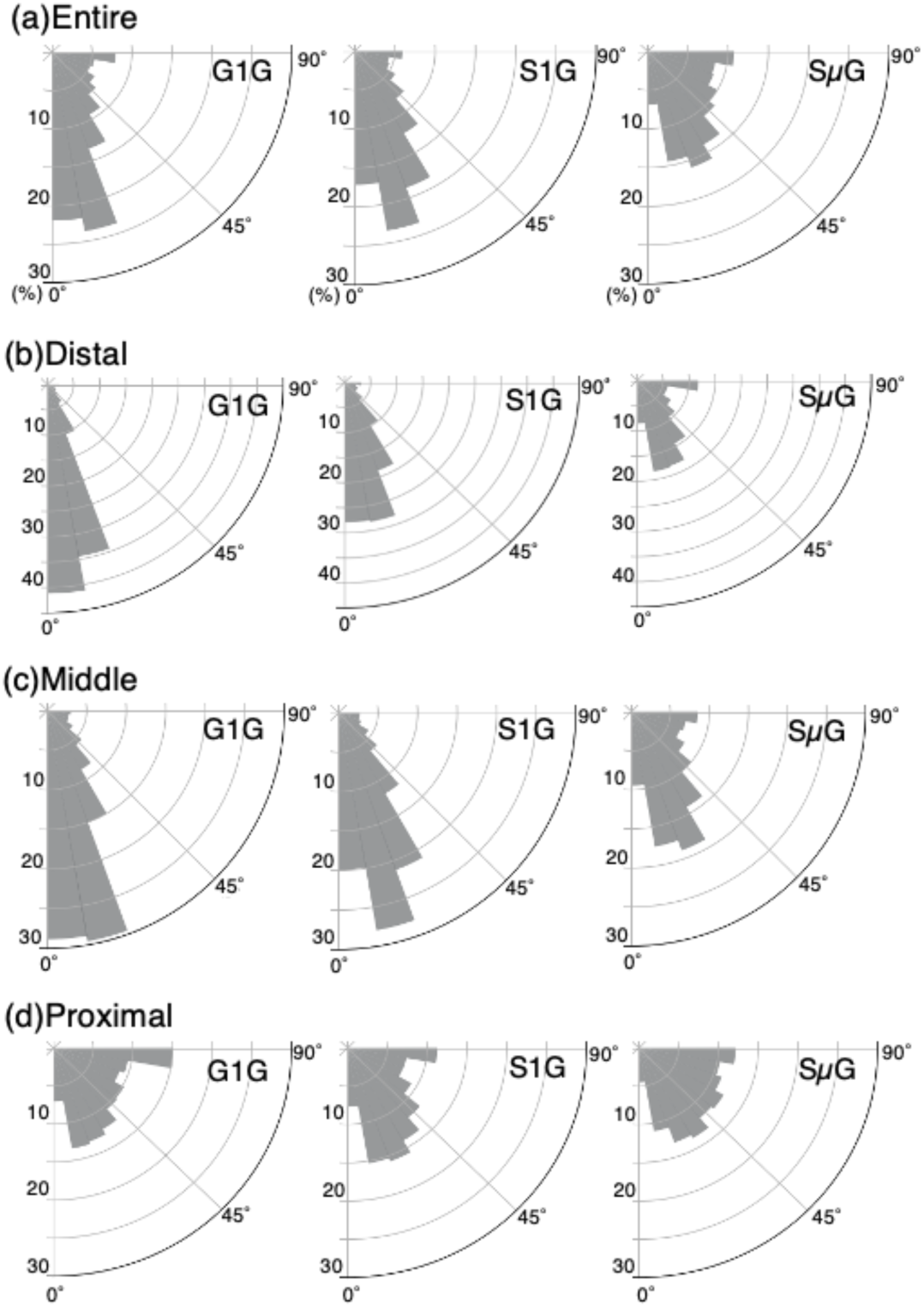
Raw angular distributions among different gravity conditions in the entire (a), proximal (b), middle (c), and distal (d) regions. Blue, red, and green columns represent Ground 1 × g, Space 1 × g, and Space µ × g, respectively. Smaller angles (closer to 0°) indicate more vertical rhizoid growth, while larger angles (closer to 90°) represent more horizontal growth orientations.

## 4. Discussion

### 4.1 Improvement of automatic rhizoid segmentation using machine learning

In this study, we improved the previous automatic rhizoid segmentation method for *P. patens* by introducing additional preprocessing steps, modifying the classifier construction and training procedure, and revising the postprocessing process. As a result, the segmentation accuracy was substantially enhanced. In addition, compared with the previous method, the present method exhibited reduced variability in prediction accuracy (Supplementary Fig. 9). This improvement is likely due to the distributed sampling of training regions along the depth axis, which enabled the classifier to better adapt to differences in rhizoid density and thus increased its generalization capability.

However, even in the present method, only two samples achieved a median F1 score above 0.8, while two others fell below 0.7. The current level of accuracy is considered to be the limit as long as the present method is used, and further improvement in accuracy would require a different approach. Moreover, in the Space µ × g samples, performance metrics remained variable among samples (Supplementary Table 1). One possible reason is that, unlike the other samples, the Space µ × g dataset exhibited large variations in object density depending on the imaging depth used to generate the ground-truth annotations (Supplementary Fig. 10). Consequently, test images differed in object density: images with dense rhizoids yielded high prediction accuracy, whereas those with sparse rhizoids showed reduced detection performance, resulting in greater variability in F1 scores. These results suggest that the classifier did not fully acquire robustness to density differences during training, leading to reduced generalization and unstable segmentation performance.

Hence, further efforts are needed to improve segmentation accuracy and enhance model generality. As an alternative to overcome the drawbacks of the current approach, convolutional neural networks (CNNs), which have begun to be applied for plant phenotyping (Brahimi et al., 2017; Shen et al., 2020; Smith et al., 2020; Vijayan et al., 2024; Xie et al., 2024; Collings and Karahara, 2026), are valid candidates. Future work will focus on implementing 3D CNN-based segmentation models and optimizing data augmentation strategies to achieve higher segmentation performance and greater automation in image analysis.

### 4.2 Three-dimensional accuracy evaluation

To examine whether the present method is applicable to morphological analysis, morphological parameters were compared between manually-created annotation volumes and the predicted volumes generated by the present automatic segmentation method. Although the results showed no significant differences between the two volumes in morphological parameters, such as, total rhizoid volume, total surface area, total rhizoid length, and length-weighted average rhizoid growth angle, the predicted volumes showed a significantly lower diameter than the ground truth (Fig. 5). This decrease in diameter was likely caused by the overlabeling of small noise particles, which reduced the average diameter. In addition, expansion of rhizoid contours was observed (Fig. 4). The contour expansion can be attributed to the use of a mean filter in postprocessing, which smoothed the boundary regions and consequently caused blurring and inflation of rhizoid outlines. Nevertheless, the presence of numerous small noise labels is suggested to largely contribute to lowering the Least Square (LS) mean diameter. To mitigate the problem of noise overlabeling, introducing CNNs, which is expected to reduce excessive noise labeling (Hendrycks et al., 2019), might be effective. Fragmentation of rhizoids was observed in predicted volumes (Fig. 4). The fragmentation is due to artifacts that occurred during sample embedment, causing the rhizoid morphology to appear in three different types (Fig. 2) and the gaps between the different types were difficult to reconnect by simply filling them. Thus, additional morphological postprocessing that considers contour connectivity will be required for correction. Since the mean filter used in the present method caused contour expansion, replacing it with a bilateral filter or anisotropic diffusion filter is considered promising for suppressing boundary blurring and inflation.

### 4.3 Differences in rhizoid elongation, volume, and surface area under different gravity conditions

In this study, the Space µ×g samples exhibited a tendency toward reduced total rhizoid volume, total rhizoid surface area, and total rhizoid length (Fig. 6). This suggests that rhizoid elongation and growth were destabilized under microgravity conditions. Generally, rhizoids in bryophytes acquire polarity and grow downward in response to gravitational stimuli (Lobachevska et al., 2022). In *P. patens*, both caulonemata and gametophores show negative orthogravitropism (Jenkins et al., 1986). Rhizoids, like protonemata, are tip-growing polarized cells. Disruption of such polarity leads to loss of local growth control, resulting in reduced elongation rates and morphological abnormalities (Hepler et al., 2001; Moody, 2022). These findings imply that the absence of gravitational cues may disturb polarity control of rhizoid, thereby reducing its elongation growth. Accordingly, under microgravity conditions, the loss of gravity stimuli likely destabilizes polarity, leading to decreased rhizoid growth. The observed reduction in rhizoid volume and surface area reflects a decrease in overall biomass, suggesting that the spatial development of the rhizoid network was restricted. Similarly, in *A. thaliana*, length of root hair, which shows tip growth, was reduced under microgravity (Kwon et al., 2015). In this case, both actin-dependent and actin-independent genetic pathways are indicated to be involved in regulation of root hair growth in space. It remains to be determined whether such processes are also involved in the reduced growth of moss rhizoids under microgravity.

Morphological analysis revealed that, the mean rhizoid diameter was significantly reduced in the Space µ × g samples (Fig. 6d). Rhizoids of *P. patens* are tip-growing polarized cells, similar to protonemata. Therefore, it is possible that the loss of gravitational cues caused irregularities in wall extension control, leading to suppression of radial expansion. However, the three-dimensional accuracy evaluation showed that the predicted volumes had a significantly lower diameter than the ground truth (Fig. 5e), whereas the predicted rhizoids tended to appear slightly thicker and exhibited expanded contours compared to the ground truth (Fig. 4b). Given these contradictions, and the fact that the mean diameter differed only slightly, despite being statistically significant, it may be premature to draw further conclusions. Improvements in segmentation accuracy will be necessary before this issue can be discussed further.

### 4.4 Changes in rhizoid growth angle under microgravity

In both the Ground 1 × g and Space 1 × g conditions, the rhizoids predominantly grew in the distal direction, as shown in the overall 3D model view (Fig. 7) and the polar histogram of raw angle values (Fig. 8). This observation indicates that the *P. patens* rhizoids show gravitropism, supporting the previous reports (Zhang et al., 2019; Lobachevska et al., 2022). Additionally, the preferential distal orientation and elongation of rhizoids suggest that this growth pattern may contribute to anchorage to the substrate against the self-weight imposed under 1 × g conditions. In the Space µ × g samples, the variance of length-weighted rhizoid growth angles was markedly larger (Table 1), indicating a reduction in the directional alignment toward the gravity vector. This suggests that, in the absence of gravitational stimuli, the rhizoids could not establish a unified growth orientation and instead elongated in random or oblique directions. In bryophyte species, such as *Ceratodon purpureus* (Hedw.) Brid., *P. patens*, *Funaria hygrometrica* Hedw., rhizoids typically exhibit positive gravitropism, whereas chloronemata and caulonemata show negative gravitropism, while caulonemata appear more sensitive to gravity and the gravi-response of rhizoids is described as weak (Lobachevska et al., 2022). Under dark conditions, caulonemal cells of *P. patens* grew in all directions when subjected to clinorotation (Jenkins et al., 1986). Similarly, in *C. purpureus*, protonemal cells grown under microgravity conditions during spaceflight completely lose gravitropism and grew in all directions (Kern et al., 2005).

Although previous studies observed growth direction on a two-dimensional plane, the present study examined growth in the three-dimensional Z-axis. Therefore, a direct comparison is not possible. However, unlike the reports under dark conditions in those earlier studies, the present study did not observe a complete loss of directional growth. Furthermore, the present study was performed under microgravity conditions with white light illumination from the upper direction. It has been reported that chloronemata show positive phototropic responses under far-red or low-intensity red light, whereas they exhibit negative phototropism under high-intensity red light (Jenkins and Cove, 1983). While phototropic responses in rhizoids have not been well characterized, the possibility of light influencing their growth cannot be excluded because the rhizoids were grown in the continuous light conditions in this experiment. Further investigation into the role of light in rhizoid development remains a subject for future study. In addition, caulonemata and chloronemata may also be present in the analyzed samples because the plants in this experiment were grown in the continuous light conditions and these structures could not be distinguished in our dataset. Although these structures are assumed to be far fewer than rhizoids, this issue requires clarification through higher-resolution observations in future studies.

In *P. patens*, the tip growth of protonemata is tightly regulated through the coordination of apical Ca²⁺ gradients and actin dynamics (Bascom Jr et al., 2018). In the case of negative gravitropism shown by protonemata, the direction of bending in response to gravitational stimulation is determined by the asymmetric distribution of actin, which depends on the minus-end–directed kinesin motor (Li et al., 2021). Therefore, it is possible that such molecular mechanisms may be weakened under microgravity, increasing variance in rhizoid growth angles.

## 5. Conclusion

Taken together, the results obtained in this study demonstrate that gravitational stimuli act as a fundamental regulatory factor in rhizoid morphogenesis, and that the absence of gravity exerts multifaceted effects on orientation and thickening growth. Under microgravity conditions in particular, increased variance in growth angles and growth inhibition were observed, resulting from the reduced of directional control. This finding suggests that, similar to seed plants, bryophytes also possess gravity-sensitive cellular response systems. In contrast, the results from the Space 1 × g samples indicate that morphological homeostasis can be restored through the application of artificial gravity. These findings provide essential baseline data for future studies in space plant physiology and morphogenesis, as well as for the development of artificial-gravity-based plant cultivation systems in space environments.

## Acknowledgements

This work was supported by the Division of Instrumental Analysis at University of Toyama. The synchrotron radiation experiments were performed at the BL20B2 of SPring-8, with the approval of the JASRI (Proposal nos. 2022B1143, 2021B1316, 2024B1190).

## Supplementary Materials

**Supplementary Table. 1.** Comparison of median F1 score, IoU, precision, and recall between the previous and present methods for each sample.

| Sample | Previous Method |  |  |  | Present Method |  |  |  |
| --- | --- | --- | --- | --- | --- | --- | --- | --- |
|  | F1 | IoU | Precision | Recall | F1 | IoU | Precision | Recall |
| G1G 27 | 0.510 | 0.343 | 0.574 | 0.474 | 0.713 | 0.554 | 0.811 | 0.624 |
| G1G 82 | 0.710 | 0.550 | 0.695 | 0.693 | 0.766 | 0.621 | 0.849 | 0.728 |
| G1G 84 | 0.476 | 0.312 | 0.548 | 0.517 | 0.792 | 0.656 | 0.864 | 0.779 |
| S1G 003 | 0.671 | 0.505 | 0.727 | 0.743 | 0.783 | 0.643 | 0.902 | 0.703 |
| S1G 004 | 0.823 | 0.699 | 0.813 | 0.866 | 0.840 | 0.723 | 0.947 | 0.779 |
| S1G 103 | 0.738 | 0.585 | 0.777 | 0.716 | 0.813 | 0.685 | 0.889 | 0.787 |
| SμG 002 | 0.133 | 0.071 | 0.409 | 0.120 | 0.637 | 0.467 | 0.688 | 0.614 |
| SμG 101 | 0.665 | 0.498 | 0.774 | 0.621 | 0.795 | 0.659 | 0.899 | 0.718 |
| SμG B101 | 0.317 | 0.189 | 0.519 | 0.360 | 0.650 | 0.482 | 0.842 | 0.539 |

**Supplementary Fig. 1.**
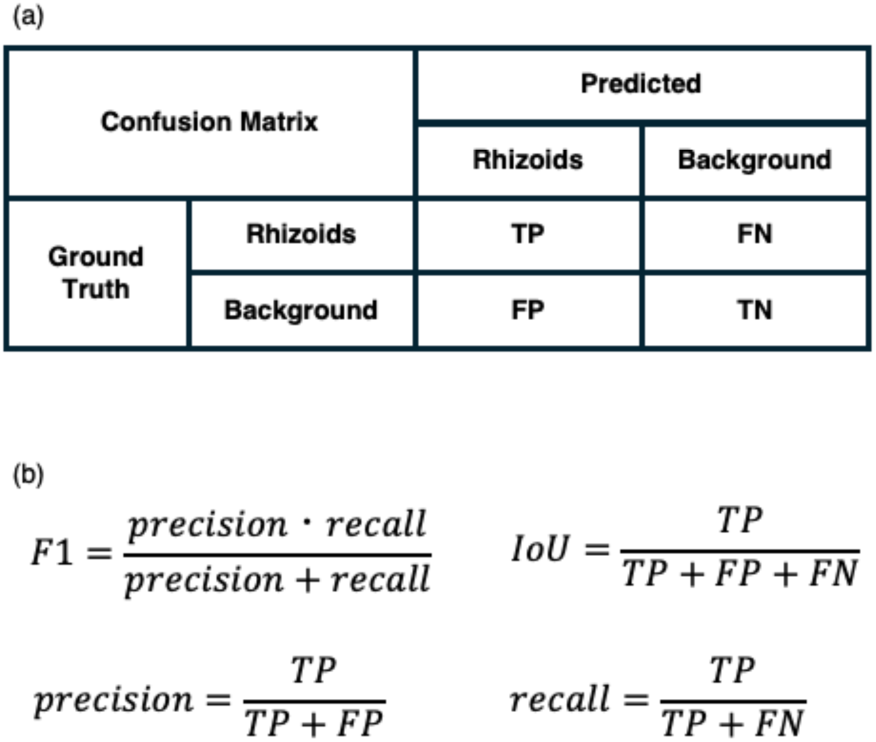
Assessment of the automatic segmentation methods. Rhizoid and background areas obtained by manual segmentation (ground truth) and automatic segmentation were compared and categorized into the areas of four groups: true positive (TP), false positive (FP), false negative (FN), and true negative (TN). (a) A confusion matrix was constructed by counting the number of pixels in each categorized area. (b) Formulas calculating pixel accuracy (PA), F1-score (F1), Intersection Over Union (IoU), precision, and recall.

**Supplementary Fig. 2.**
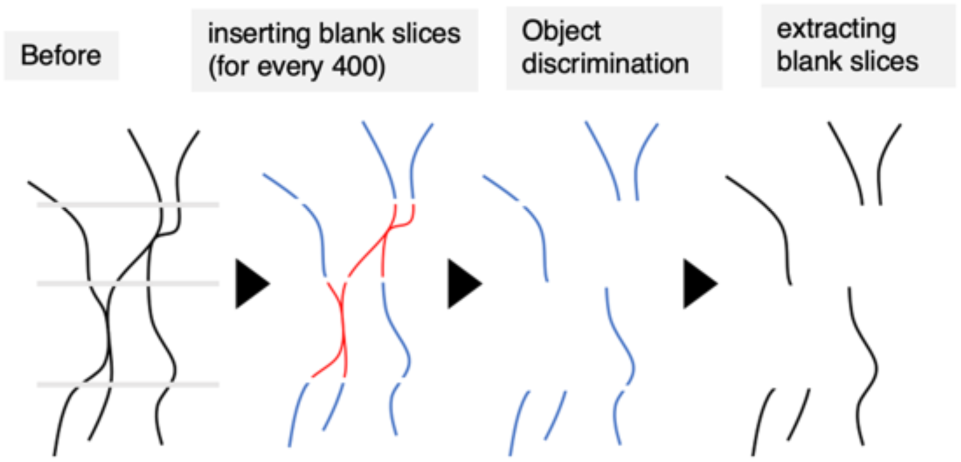
Schematic illustration depicting artificial splitting of rhizoids for object discrimination in order to quantify rhizoid growth angle.

**Supplementary Fig. 3.**
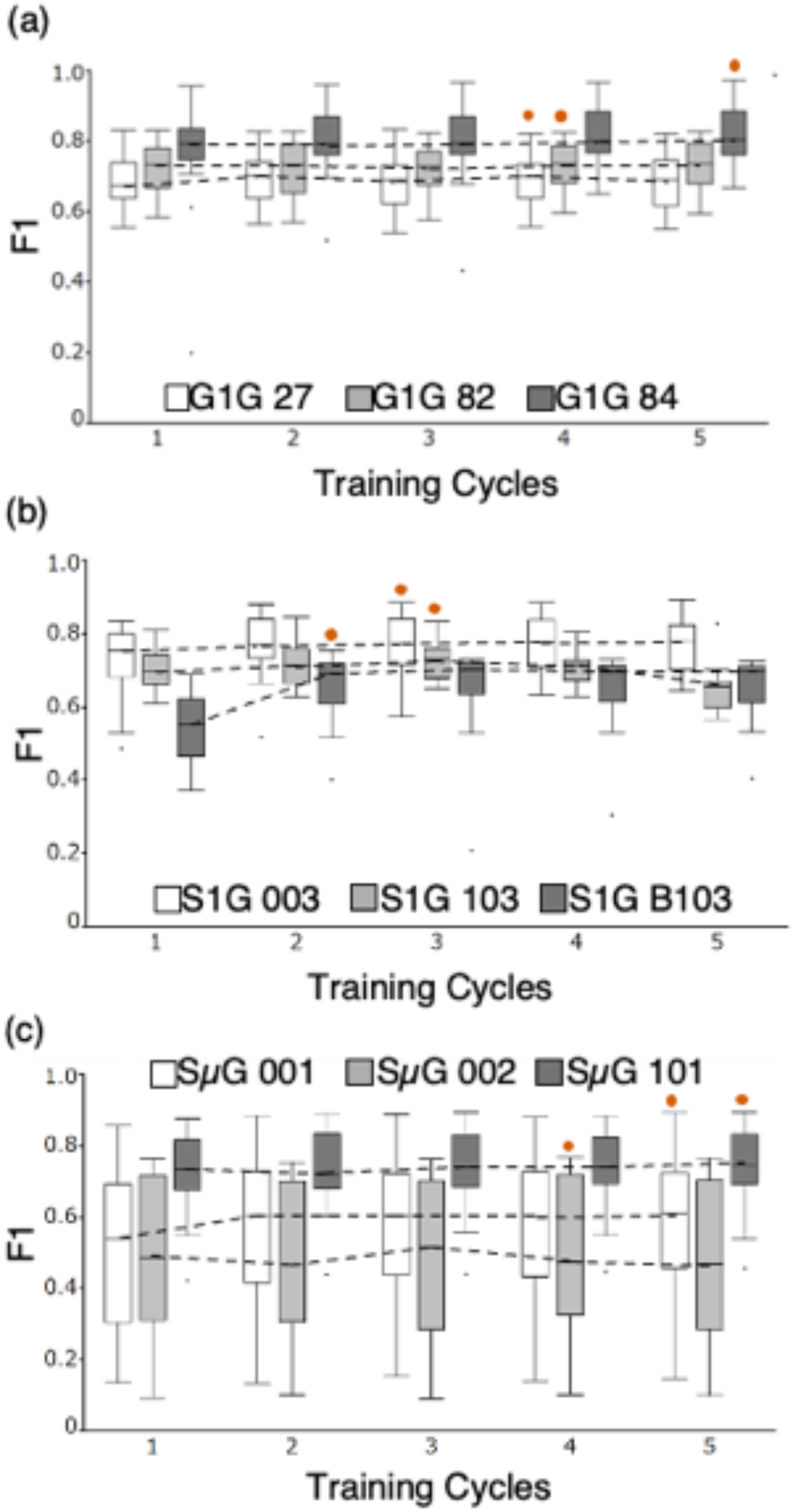
Changes in F1 scores by changing number of training cycles in Ground 1 × g (a), Space 1 × g (b), Space µ × g (c) (n = 3). Box plots display maximum, minimum, median, and first/third quartiles. Red circles within the graphs indicate the number of cycles yielding the highest average F1 score for each sample. No significant difference was detected within the same sample (Kruskal-Wallis test, *P* ≥ 0.05).

**Supplementary Fig. 4.**
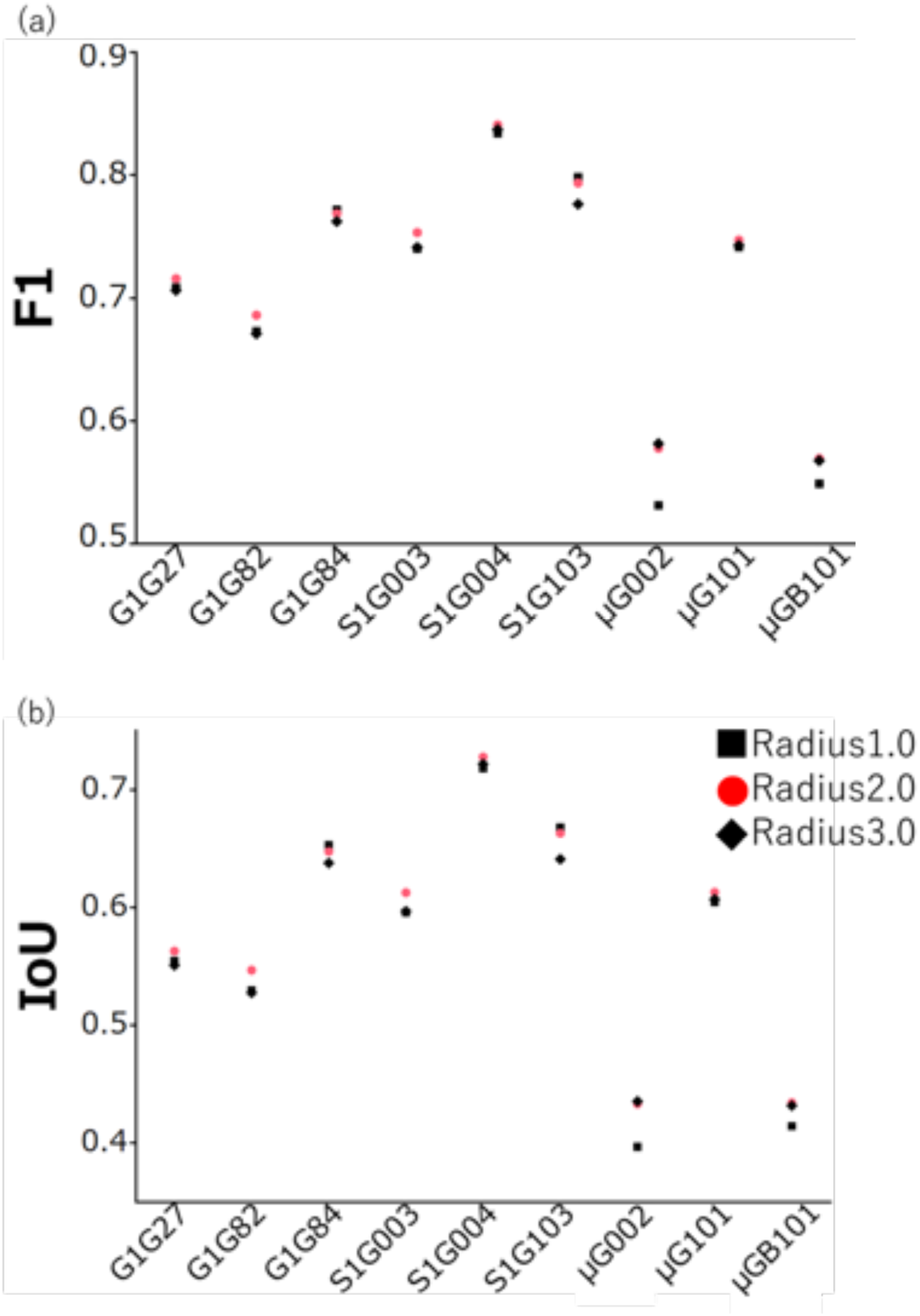
Changes in F1 scores and IoU of the samples by changing Closing Radius from 1 to 3. (a) F1 scores and (b) IoU for each sample. Twenty-seven ground truth and predicted images were used for the analysis.

**Supplementary Fig. 5.**
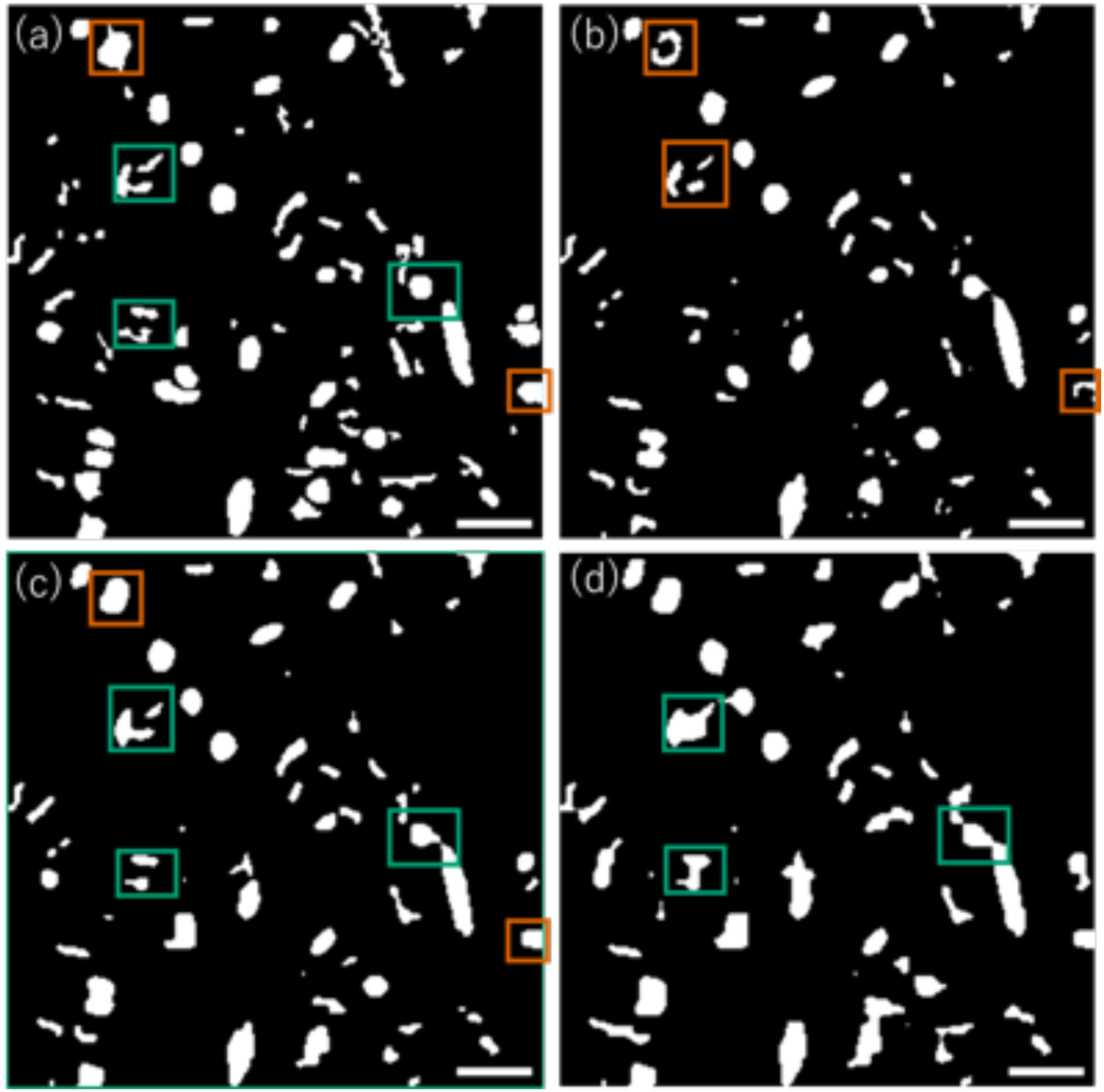
Differences in segmentation results by changing Closing Radius from 1 to 3. X-Y images from Space 1 × g_003. (a) Ground truth image, (b) Closing 1, (c) Closing 2, (d) Closing 3. The orange box highlights notable differences between Closing 1 and 2, the green one between Closing 2 and 3, and the purple one among all three conditions (1, 2, and 3). Scale bars = 0.1 mm.

**Supplementary Fig. 6.**
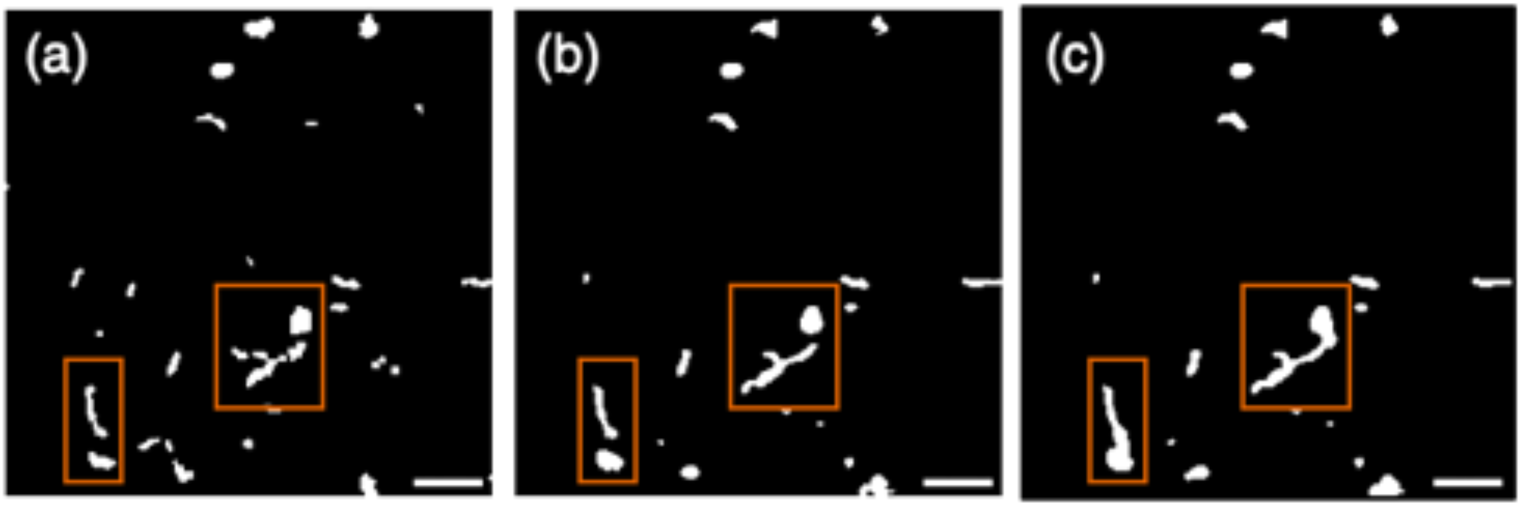
Differences in segmentation results between Ground truth (a), Closing 1 (b), and 2 (c) in X-Y images of Ground 1 × g_84, where Closing 1 achieved the highest performance. Orange box highlights areas with marked differences due to Closing parameter changes. Scale bars = 0.1 mm.

**Supplementary Fig. 7.**
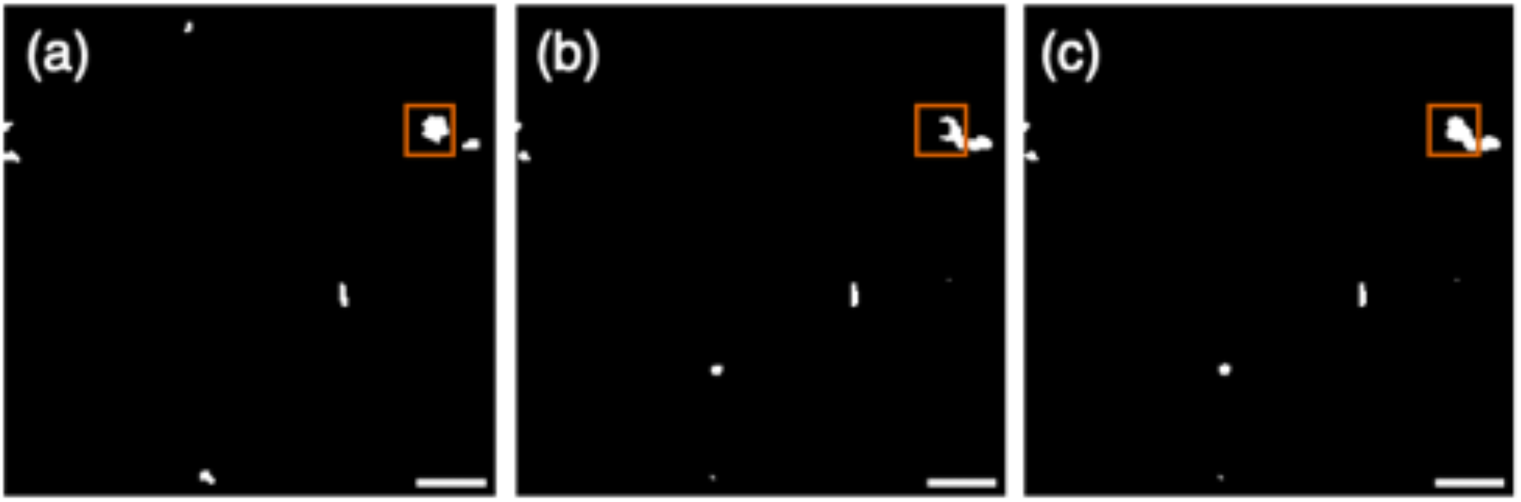
Differences in segmentation results between Ground truth (a), Closing 2 (b), and 3 (c) in X-Y images of Space µ × g_002, where Closing 3 achieved the highest performance. Scale bars = 0.1 mm.

**Supplementary Fig. 8.**
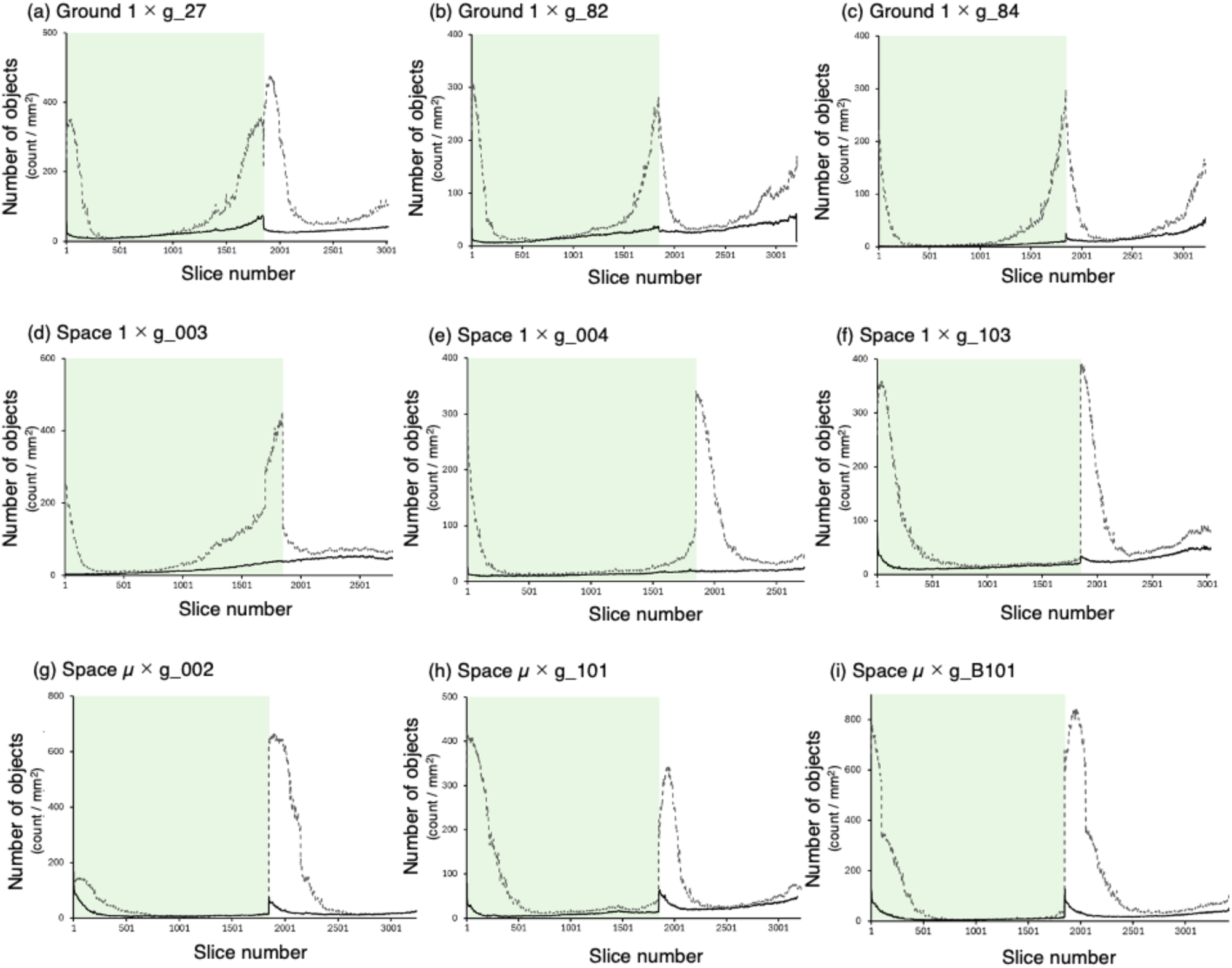
Changes in the distribution of the objects along the depth of the volume methods in Ground 1 × g_27 (a), Ground 1 × g_82 (b), Ground 1 × g_84 (c), Space 1 × g_003 (d), Space 1 × g_103 (e), Space 1 × g_004 (f), Space µ × g_002 (g), Space µ × g_101 (h), and Space µ × g_B101 (i). Broken line, before preprocessing. Solid line, after postprocessing.

**Supplementary Fig. 9.**
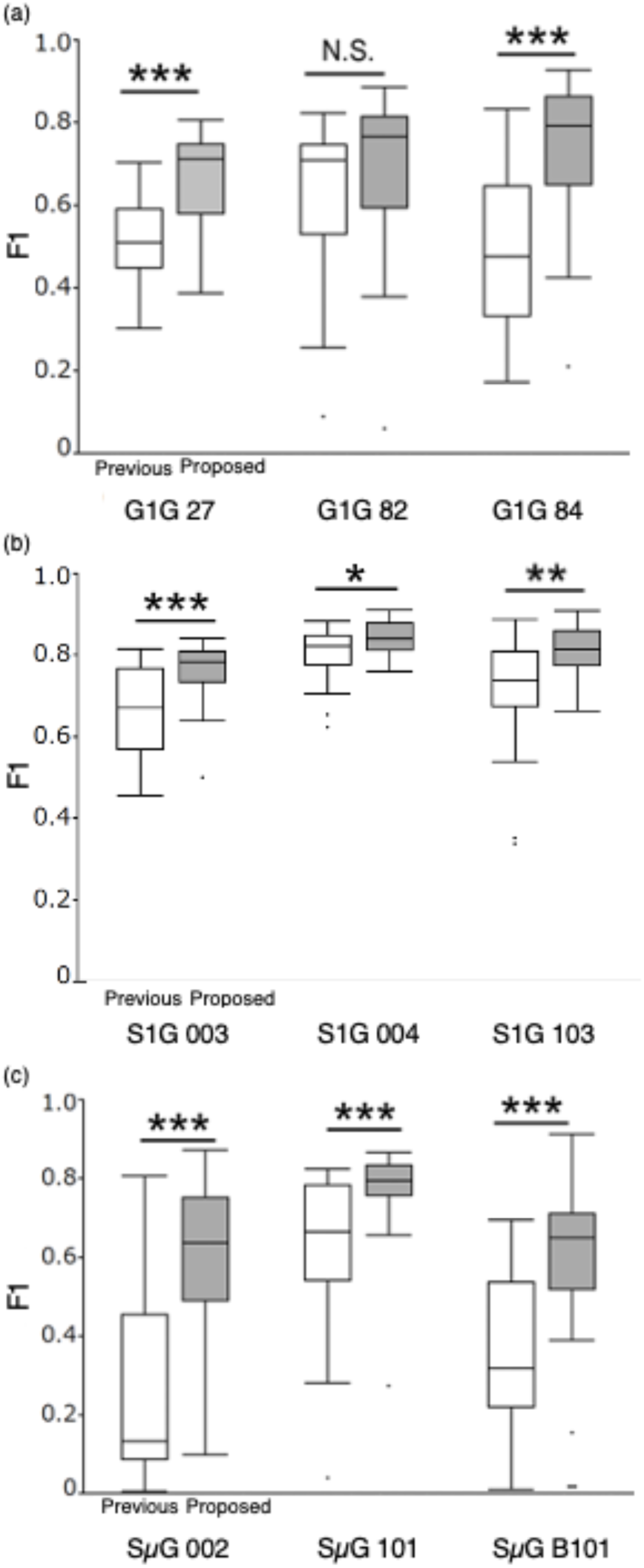
Comparison of F1 scores of the samples between the previous and the present methods in Ground 1 × g (a), Space 1 × g (b), and Space µ × g (c). Twenty-seven sets of ground truth and predicted X-Y images were used for the analysis. Box plots display the F1 scores of the previous (open) and the present method (shaded). Mann-Whitney *U*-test (two-tailed) was used. N.S., not significant; *, *P* < 0.05; **, *P* < 0.01; ***, *P* < 0.001.

**Supplementary Fig. 10.**
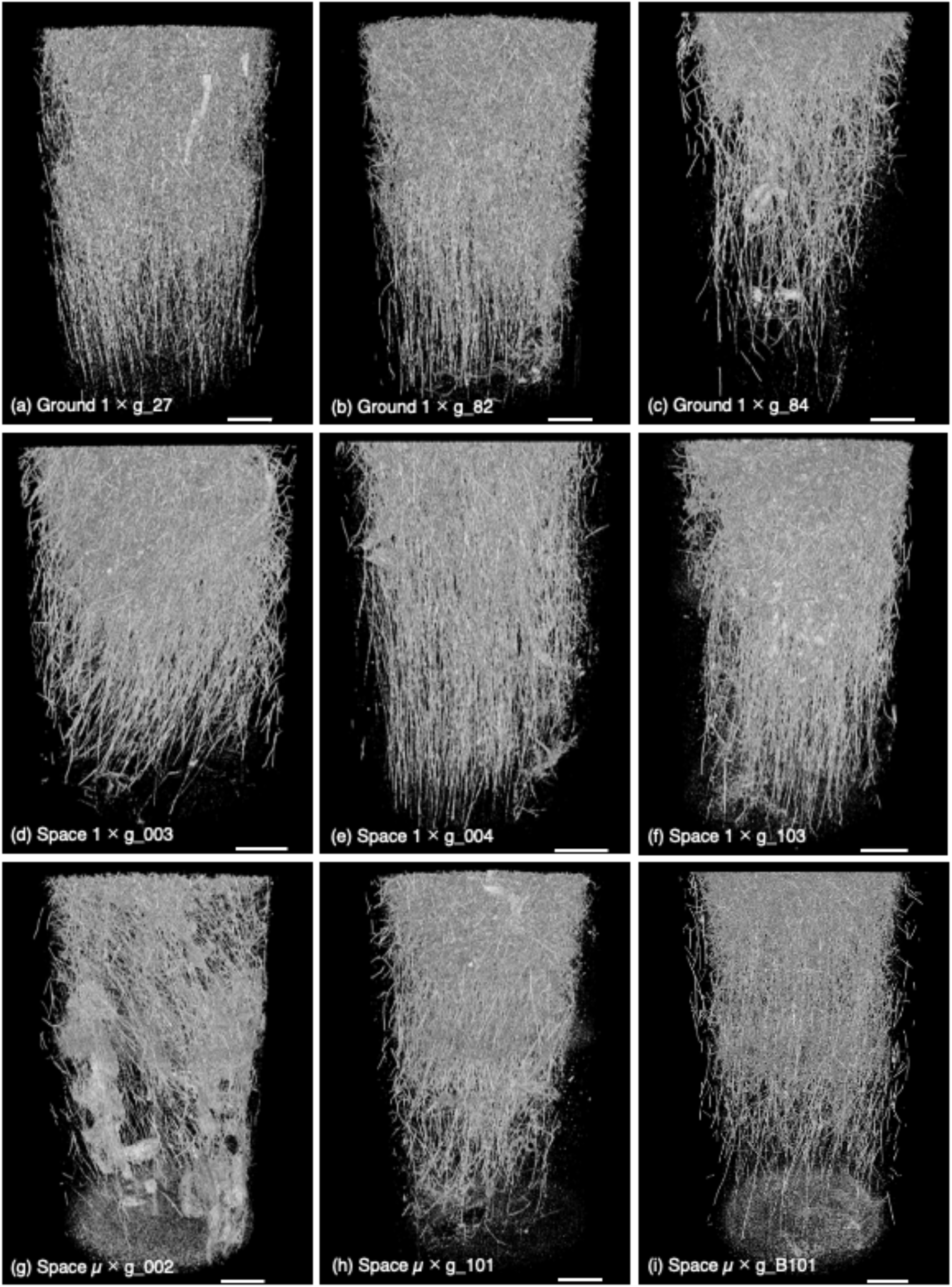
3D models of the rhizoid system architecture in Ground 1 × g_27 (a), Ground 1 × g_82 (b), Ground 1 × g_84 (c), Space 1 × g_003 (d), Space 1 × g_004 (e), Space 1 × g_103 (f), Space µ × g_002 (g), Space µ × g_101 (h), and Space µ × g_B101 (i).

